# BooLEVARD: Boolean Logical Evaluation of Activation and Repression in Directed pathways

**DOI:** 10.1101/2025.03.24.644921

**Authors:** Marco Fariñas, Eirini Tsirvouli, John Zobolas, Tero Aittokallio, Åsmund Flobak, Kaisa Lehti

## Abstract

Boolean models are widely used for studying dynamic processes of biological systems. However, their inherent discrete nature limits their ability to capture continuous aspects of signal transduction, such as signal strength or protein activation levels. Although existing tools provide some path exploration capabilities that can be used to explore signal transduction circuits, the computational workload often requires simplifying assumptions that compromise the accuracy of the analysis. Here, we introduce BooLEVARD, a Python package designed to efficiently quantify the number of paths leading either to node activation or repression in Boolean models, which offers a more detailed and quantitative perspective on how molecular signals propagate through signaling networks. By focusing on the collection of non-redundant paths directly influencing Boolean outcomes, BooLEVARD enhances the precision of signal strength representation. We showcase the application of BooLEVARD in studying cell-fate decisions using a Boolean model of cancer metastasis, demonstrating its ability to identify critical signaling events. Furthermore, BooLEVARD provides a refined understanding of signaling dynamics, which can increase our understanding of disease development and drug responses. BooLEVARD is freely available at https://github.com/farinasm/boolevard.

## Introduction

The rapid growth of high-throughput technologies has enabled the generation of vast amounts of biological data, leading to a critical challenge of its analysis and interpretation (1). Computational modeling methods have provided enhanced understanding of cellular processes, supporting experimental research. One example is agent-based modeling of individual components of a system, known as agents, dynamically interacting according to predefined rules (2). These rules can be based on mathematical frameworks, such as Boolean equations, constraint-based modeling (CBM), or differential equations (3–5). While differential equations are particularly useful for modelling continuous processes, such as biochemical reactions, molecular transport, or diffusion, their use is often limited by the requirement for detailed kinetic experimental data, which are typically hard to acquire (6). Boolean equations and CBMs are more easily applicable to discrete processes. Even though CBMs, like genome-scale metabolic modeling, also require certain parameters, such as omics data and culture media composition, these are easier to obtain (3). Boolean models are typically based on signaling diagrams representing events such as transcriptional regulation and protein-protein interactions. Boolean models do not require any kinetic data and can be constructed based on causal interactions (4). As such, Boolean modeling provides a simplified representation of dynamic systems, while still effectively capturing the key mechanisms that govern biological processes and lead to cellular phenotypes (7). Moreover, their less stringent information requirements make them a practical choice for modeling a wide range of biological systems.

Boolean models consist of a network of interacting nodes, each representing a biological entity that can exist in one of two states: active (i.e. 1) or inactive (i.e. 0) (4). The state of a node, referred to as its Boolean or local state, is governed by Boolean equations that define how the states of upstream regulators are integrated, and depend on the network’s topology (8). During the model simulation, the user can specify an initial condition (i.e., a set of active and inactive nodes), allowing the model to explore all possible state transitions. States that have no outgoing transitions are known as attractors, which can be further classified into stable states or cyclic attractors, and are expressed as Boolean vectors of local states that frequently can represent cellular phenotypes (9). Attractor reachability, starting from given initial conditions until the attractors are reached, can be explored with several computational tools available for analyzing Boolean models. Examples include bioLQM, mpbn, Pint, and GINsim (10–14). Other tools, such as Ma-BoSS and PhysiBoSS (15–17), extend the Boolean modeling by incorporating continuous Markov chains and allow for the study of a system’s evolution over time, and explore the probabilities of node activities and stable state reachability (15).

Despite their advantages, Boolean models inherently provide only binary information (whether a node is active or inactive), without capturing the intensity of activation or inhibition (18). To address this limitation, we introduce BooLEVARD (Boolean Logical Evaluation of Activation and Repression in Directed pathways), a Python package that leverages equations within Boolean models to systematically count the number of activating and repressing paths leading to the local state of a given node. Unlike existing approaches, which typically do not quantify repression and often require users to impose a maximum path length constraint to manage computational complexity (12, 19), BooLEVARD enables the exploration of activatory and inhibitory signal intensities. In the present study, we demonstrate how signal transduction strength influences invasive and apoptotic fate decisions using a Boolean model of cancer metastasis (20). Moreover, perturbation performance and downstream analysis with BooLEVARD allowed for a more detailed and granular distribution of the perturbation impact. Overall, this novel approach provided a more insightful interpretation, compared to traditional stable state analysis, therefore expanding the capabilities offered by Boolean models to understand the underlying complexity of biological systems.

## Design and Implementation

### Requirements

BooLEVARD is available in Python 3 as a PyPi package, providing an efficient and user-friendly interface that can be implemented in Jupyter Notebooks. Detailed installation instructions and a quickstart tutorial, are available in the package’s GitHub repository (https://github.com/farinasm/boolevard).

### Inputs and the BooLEV class

BooLEVARD takes Boolean models in BoolNet format (.*bnet*) as input (21), which are loaded and managed as objects of the BooLEV class. Each BooLEV object provides structured access to key model information, including a list of nodes, a dictionary storing logical rules in canonical Disjunctive Normal Form (cDNF) (22), a dictionary for the negated rules in cDNF (referred to as cNDNF for consistency), a pandas data frame containing the model’s stable states, and a data frame that integrates all this information. This structure enables efficient exploration and downstream analysis of Boolean models within BooLEVARD. The stable states of the model are automatically computed by the Most Permissive Boolean Network (mpbn) package (11), while the cDNFs and cNDNFs are determined with the Python Electronic Design Automation (PyEDA) package (23). BooLEV objects implement four methods: *CountPaths, Drivers, Pert*, and *Export*. The workflow of BooLEVARD is illustrated in **Figure 1**.

**Figure 1.**
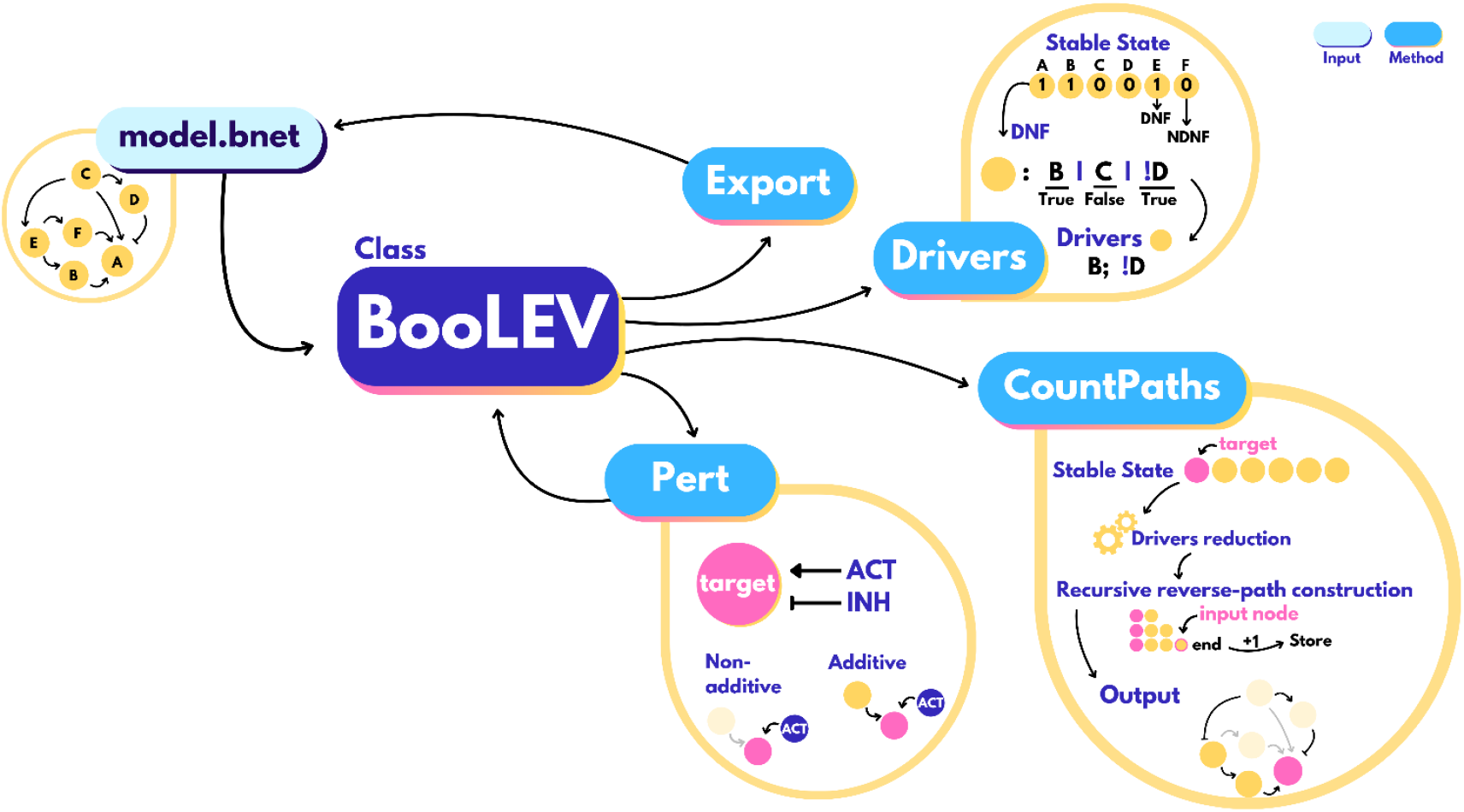
Overview of the BooLEVARD workflow. BooLEVARD processes Boolean models in .*bnet* format by converting them into *BooLEV* objects, integrating the *Export, CountPaths, Drivers*, and *Pert* methods. The *Export* method allows users to save *BooLEV* objects back as .*bnet* files, ensuring compatibility with other tools. The *CountPaths* method performs BooLEVARD’s core function – counting the number of activatory and inhibitory paths of a specific node within the stable states of the model. Stable-state-specific *drivers* (conjunctive blocks within a Boolean equation in cDNF) of each node are calculated using their cDNFs (for active nodes) or cNDNFs (for inactive nodes). *Drivers* can also be computed by calling the *Drivers* method. The *drivers* dictionary is simplified by applying iterative transitive reductions, and paths are recursively reversed-constructed and stored in a counter, being this the output of the *CountPaths* method. The *Pert* method enables users to introduce perturbations into *BooLEV* objects by modifying the Boolean equations of target nodes. Perturbations can be either additive, where the perturbation is incorporated into the existing logical rule, or non-additive, where the original equation is completely substituted to exclusively reflect the perturbation’s effect. The updated *BooLEV* can be exported back to .*bnet* format.

### Recursive reverse-path construction and enumeration

BooLEVARD’s main feature is to count the number of paths leading to the activation or inactivation of a specific node within a stable state reached by a Boolean model (both selected by the user), achieved using the *CountPaths* method. To determine the number of paths, BooLEVARD employs a recursive reversed-path construction strategy, building paths from the target back to the input nodes of the network (i.e. those with no incoming regulation). The first step in this process involves checking the Boolean state of each node in the chosen stable state to retrieve either their cDNFs (for active nodes) or the cNDNFs (for inactive nodes). The cDNF provides a non-redundant, maximally simplified representation of a Boolean equation as a disjunction of conjunctions, where each conjunctive block represents a potential step in the transduction pathway leading to the node’s Boolean state (22). These blocks serve as the fundamental components of the paths analyzed by BooLEVARD. The *CountPaths* method then uses the c(N)DNFs to compute a dictionary of drivers. Drivers are defined as stable-state-specific conjunctive blocks (within cDNFs and cNDNFs) driving the Boolean state of the rule’s target node. Accordingly, the Boolean outcome of a driver is always true. This computation is independently available through the *Drivers* method of BooLEV objects. Once the drivers are computed, a simplification process is applied through iterative transitive reductions (24) across the dictionary, where redundant connections are removed while conserving the signal transduction dynamics. This simplification step notably reduces the overall complexity and eases the computational workload in successive steps. Following this, BooLEVARD uses the processed drivers of the target node each being the next upstream unit of a potential path (**Figure 1**).

The resulting set of partial paths is analyzed to ensure that none of the paths are repeated; any duplicates are discarded. This process is applied recursively, continuing until all paths are fully processed – that is, until they either form a loop or reach an input node, a condition that is checked at every elongation step. Accordingly, the tool identifies only non-loop paths, which are defined as linear or complex. Linear paths (**Figure 2A**) are built when single nodes are added at each stage, while complex paths (**Figure 2C**) arise when a step involves multiple nodes linked by a conjunction. The difference between complex paths and loops (**Figure 2B**) is that when a conjunction appears, at least one of the nodes involved reaches an input (i.e. results in a linear path). If a partial or total linearity is not fulfilled, the path elongation stops, and the path is discarded. Finally, each valid path (i.e. linear path reaching the target from an input node) is added to a counter, being the final count (positive or negative, depending on whether the local state of the target node is 1 or 0, respectively) which is the output of the *CountPaths* method. Since the number of valid paths represents the total number of independent ways in which a node can be activated or repressed, a higher path count indicates stronger signal transduction toward the target node, providing a measure of signaling strength.

**Figure 2.**
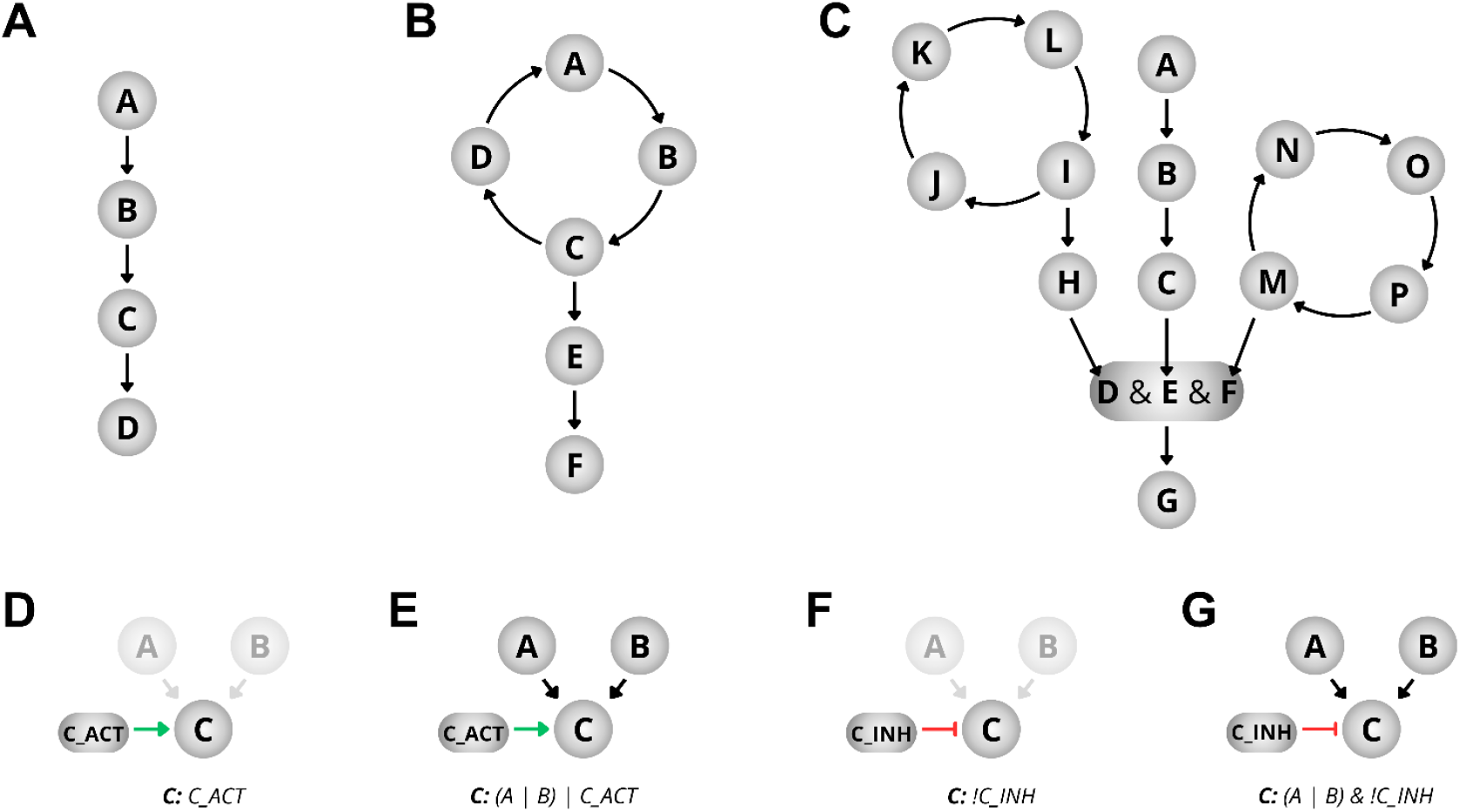
**A**. Example of a straightforward linear path. **B**. Representation of a positive feedback loop. **C**. Illustration of a complex signaling pathway. **D**. Schematic of a non-additive activation. **E**. Schematic of an additive activation. **F**. Schematic of a non-additive inhibition. **G**. Schematic of an additive inhibition.

### Model Perturbations

Tools for processing Boolean models often include functions for simulating perturbations, typically node knock-outs (KOs) or ectopic expressions (EEs) (10 ,11, 13). These perturbations are usually implemented by replacing the target node’s Boolean equation with either 0 (for KOs) or 1 (for EEs). However, since BooLEVARD constructs transduction paths directly from the model’s Boolean equations, this replacement approach is not feasible. To address this, BooLEV objects include the *Pert* method, which extends the range of perturbations from two to four types: two forms of activation and two forms of inhibition, each categorized as either additive or non-additive. Upon non-additive perturbations, the target node’s Boolean equation is substituted by a Boolean formalism that reflects only the regulation triggered by the perturbation. Additive perturbations determine the state of the target node similarly, but the effects of the perturbation are incorporated into the target node’s original Boolean equation.

To implement the perturbations, BooLEVARD creates a dedicated perturbation node that either positively (activations) or negatively (inhibitions) influences the target node. The Boolean equation of the target node is then updated to account for the user-defined additive or non-additive perturbation (**Figure 2D-G**). The resulting perturbed version of the Boolean model is stored as a BooLEV object.

## Results

The following case study illustrates how BooLEVARD enables a more granular approach to study signal transduction in Boolean models by counting the paths influencing the Boolean state of a node. Additionally, it demonstrates the importance of considering the signal strength in understanding biological processes such as cell fate decisions. We demonstrate how BooLEVARD offers increased output resolution compared to traditional stable state analysis. In this use case, we used a Boolean model of metastasis developed by Cohen *et al*., 2015 (20), which incorporates key signaling cascades driving cell invasion in cancer.

### BooLEVARD reveals variability in signal transduction strength influencing cell-fate decisions

The metastatic model (20) captures metastasis-associated and ECM- and DNA-damage-induced cell functions. Accordingly, it includes two input nodes (DNAdamage, ECMicroenv) and six phenotype nodes (Invasion, Migration, CellCycleArrest, Apoptosis, Metastasis, and EMT). The Metastasis node was not included in the analysis, as its activation is exclusively dependent on the Migration node. In addition, the TGFbeta node was split into two nodes, namely TGFbeta_i (activated by NICD and inhibited by CTNNB1) and TGFbeta_e (activated by ECMicroenv) to differentiate between intrinsic- and extrinsic-TGFβ-related regulation, respectively. Both TGFbeta_i and TGFbeta_e converge into a generic TGFbeta node.

The model produces nine stable states, including one homeostatic state (*HS*, all phenotype nodes are active), four apoptotic states (*Apop1-4, Apoptosis* is active), two EMT states (*EMT1/2, EMT* is active), and two metastatic states (*M1/2*, Invasion and Migration are active). Although stable states within the same phenotypic category share overlapping local states of phenotype nodes, analysis using BooLEVARD provided greater resolution (**Figure 3**). A clear example relies on the apoptotic stable states, which can be classified into two groups, integrated by *Apop1/2* and *Apop3/4*, respectively. *Apop1/2* are triggered upon presence of *DNAdamage* and absence of *ECMicroenv*, whereas *Apop3/4* are reached upon presence of both input nodes. Although all four states display identical cell fates, as indicated by the Boolean state of the phenotype nodes, BooLEVARD provides additional insight. By counting activatory and inhibitory paths, it predicts reduced apoptotic signaling upon presence of *ECMicroenv*. The same applies when comparing the *EMT1/2* states, where stronger EMT commitment is observed upon communication with the extracellular microenvironment. These results align well with already published studies stating cancer-ECM crosstalk confers protection against apoptosis (25). Moreover, cancer cells can undergo EMT in response to DNA damage, thus displaying increased DNA-repair potential. On the other hand, accumulation of genomic damage has been shown to have the opposite effect and trigger apoptosis (26, 27).

**Figure 3.**
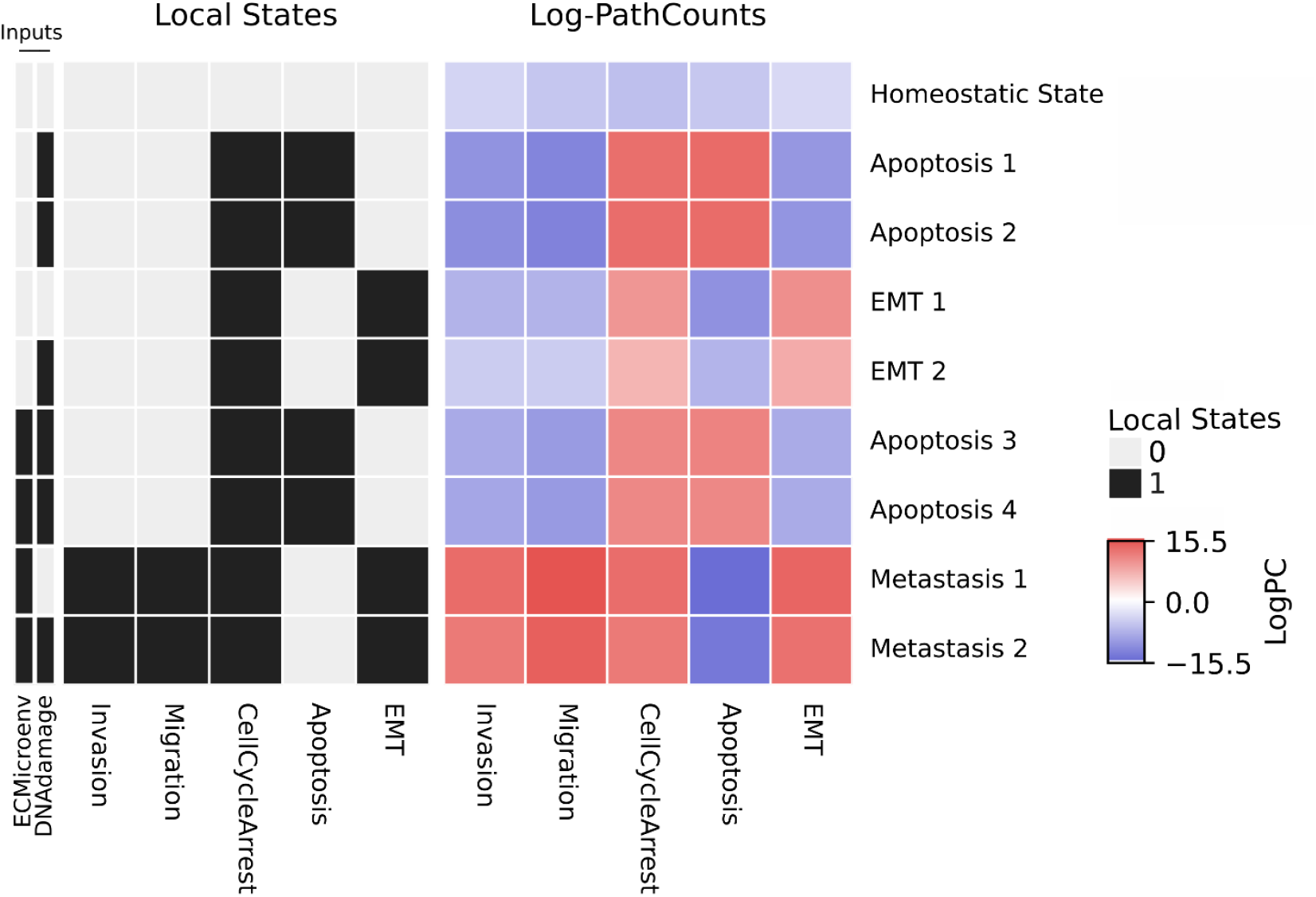
Heatmap showing the Boolean states (left) and the corresponding path counts (right), calculated using BooLEVARD, leading these states for the phenotype nodes in the stable states reached by the metastatic model. The Boolean states of the *DNAdamage* and *ECMicroenv* input nodes driving each stable state are also displayed. Path counts are presented in logarithmic scale. Active and inactive nodes are presented in black and white, respectively. Active paths triggering node activation and inactivation are presented in red in blue, respectively.

### BooLEVARD highlights negative correlation between invasion and apoptosis

To investigate whether perturbations with seemingly similar stable states, and therefore phenotypic out-comes, differ in their underlying signaling circuits, we individually perturbed each non-input and non-phenotype node in the model by simulating both additive activation and inhibition scenarios (see *Model Perturbations*). Accordingly, this approach simulates the effects of drug or other types of treatments either activating or inhibiting the proteins represented in the model. Using BooLEVARD, we evaluated the impact of these perturbations on metastasis by focusing on the *Invasion* and *Apoptosis* phenotype nodes as readouts. A total of 195 stable states were analyzed across 48 perturbations, all performed on a local computer with 16 GB RAM and AMD Ryzen 75800H processor in approximately 21 minutes (**Figure S1**).

Results were then compared to the Boolean states of *Invasion* and *Apoptosis* nodes to identify any differences both stable-state-wise and model-wise (i.e. averaging node Boolean states and path counts of the stable states of a given model). The metastatic states, *M1* and *M2*, are only reached when the *ECMicroenv* input node is activated, which also triggers two apoptotic states. Consequently, the effect of these perturbations on metastatic fate was analyzed under this initial condition. As expected, implementation of BooLEVARD resulted in high granularity across the path count distribution displayed by the perturbations (**Figure 4A-D, Supplementary Figures 2-4**). BooLEVARD allowed for tracing the intensities through which signals are transduced towards the Boolean state of a given node. As a result, differences arise although two nodes share the same Boolean state upon different perturbations, which directly highlights differences between the internal circuits and signal transduction strengths. For example, while perturbations resulting in highest fate transitions modulating invasion and apoptosis (**Figure 4E-P, Supplementary Figure S5**) show high agreement between both approaches, BooLEVARD still enables us to observe distinctions in the intensity of these effects.

**Figure 4.**
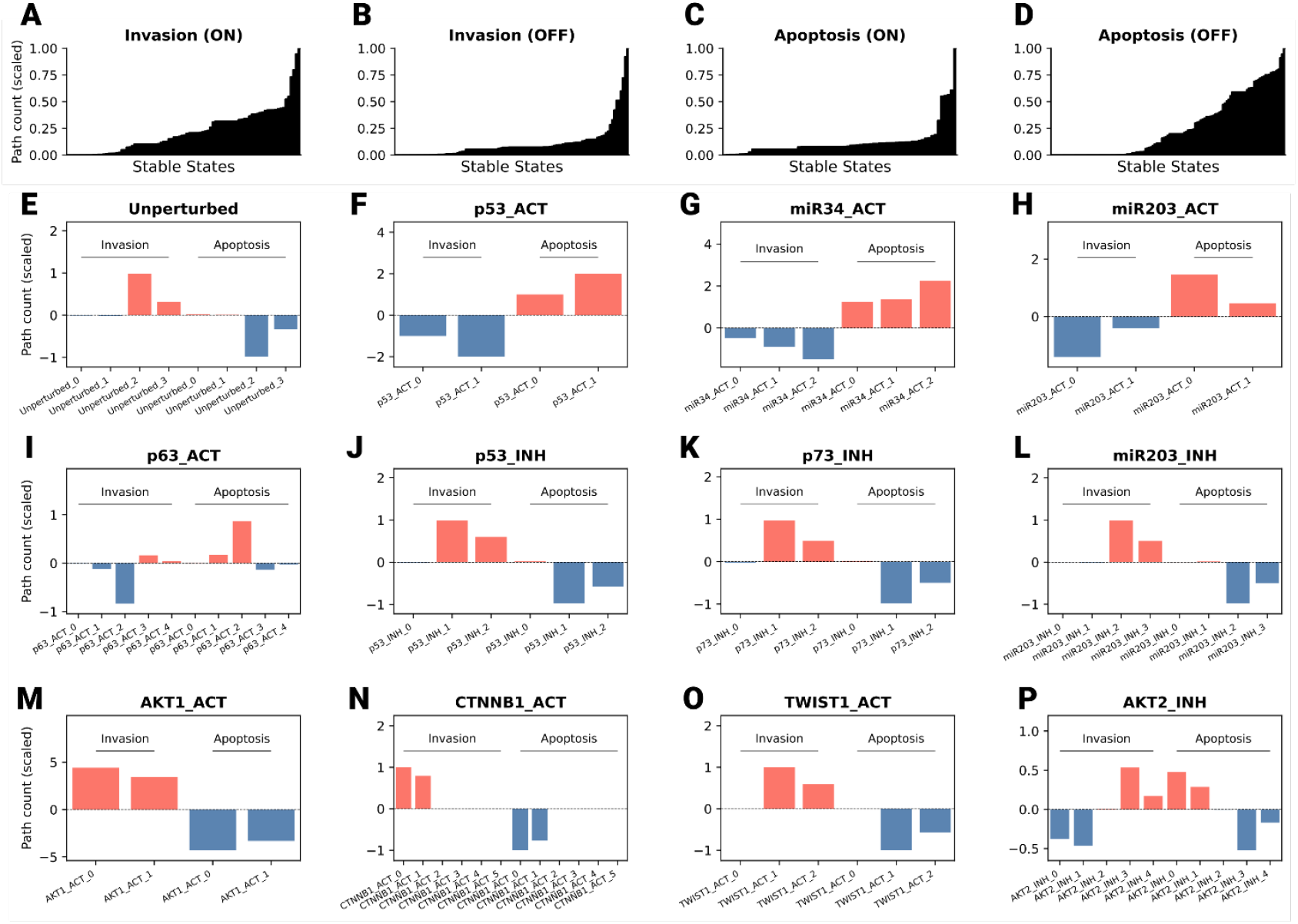
**A-D**. Path count range leading to the Boolean states of the *Invasion* and *Apoptosis* phenotype nodes across the stable states reached under the different perturbations. Path counts are shown as min-max scaled values across all stable states. **E-P**. Barplots of path counts triggering the activation and inactivation of *Invasion* and *Apoptosis* phenotype nodes across the stable states reached upon perturbation. Each bar shows the min-maxed scaled (with reference at zero) path counts in a given stable state. Activatory (red) and inhibitory (blue) paths are displayed for each target node (*Invasion* and *Apoptosis*) in different perturbations.

Notably, the stable state analysis highlights *p53_ACT, miR34_ACT*, and *miR203_ACT* (**Figure 4F-H**) as, on average, the most impactful perturbations in dampening invasion and triggering apoptosis, all three with qualitatively identical effects (**Supplementary Figure S2**). Together with *p63_ACT* (**Figure 4I**), these perturbations are displayed as the strongest triggering anti-invasive fate when using BooLEVARD. Therefore, despite overlapping invasive and apoptotic fate trends, an increasing gradient is observed in *miR203_ACT, p53_ACT*, and *miR34_ACT*, respectively. Consistent with the lowest values observed for apoptosis resistance and invasion, miR-34 has been demonstrated to enhance p53-mediated responses and to inhibit proliferation and invasion *in vivo* (28). p63, a gene related to p53, is often dysregulated in cancer and overexpression of its transactivation-domain-containing isoform (TAp63) has been linked to enhanced apoptosis and reduced metastatic potential (29) and, together with p73, it cooperates with p53 to trigger its function (30). Conversely, stable states reached upon inhibitions of p53, p73 and miR203 cell cycle regulators, and as supported by experimental evidence (31), activations of AKT1 and EMT-related genes such as CTNNB1 and TWIST1 (**Figure 4J-O**) were predicted to have the strongest impact on promoting invasion and inhibiting apoptosis.

Although not frequent, some mismatches between the two approaches were observed. An interesting case is *AKT2_INH*. When this perturbation is applied, an additional invasive stable state is reached when compared to the unperturbed setup (**Figure 4E**), therefore leading to a total of three pro-invasive and two pro-apoptotic stable states. On average, Boolean states of phenotype nodes favor invasiveness overall, but analysis with BooLEVARD showed that the two pro-apoptotic stable states exhibit notably more signaling intensity than their three pro-invasive counterparts (**Figure 4P**). These results are further supported by experimental evidence, which demonstrates that Akt2 inhibition has a negative impact in invasion and colony formation in colorectal cancer, whereas Akt1 inhibition had no effects (32). Moreover, BooLEVARD also showed that the inhibition of AKT1 resulted in little to no effect. Interestingly, BooLEVARD predicts *AKT1_ACT* to have a stronger proinvasive effect than *AKT2_ACT* (the anti-apoptotic effects of both perturbations remain similar), although stable states analysis results in overlapping effects. Akt proteins regulate key cellular processes, including protein synthesis, proliferation, invasion, and inflammation (33). In cancer, Akt1 over-activation has been linked to tumor growth during earlier invasion stages, whereas Akt2 has been shown to be more involved in facilitating distant metastases (34).

Finally, to investigate how the invasion and apoptosis variables relate to one another using both methods, we investigated the score distributions obtained from the stable states and BooLEVARD analyses, respectively (**Figure 5, Supplementary Figure S6**). Although a correlation between invasion and apoptosis (represented as apoptosis resistance in the figure) can already be observed with the stable state analysis, stronger Pearson correlation was displayed by BooLEVARD’s results (r = 0.60, p = 5.90e-6 vs. 0.77, p = 1.26-10). This further emphasized the increased granularity of BooLEVARD’s output and its value in producing results that contribute to improved interpretation of biological observations.

**Figure 5.**
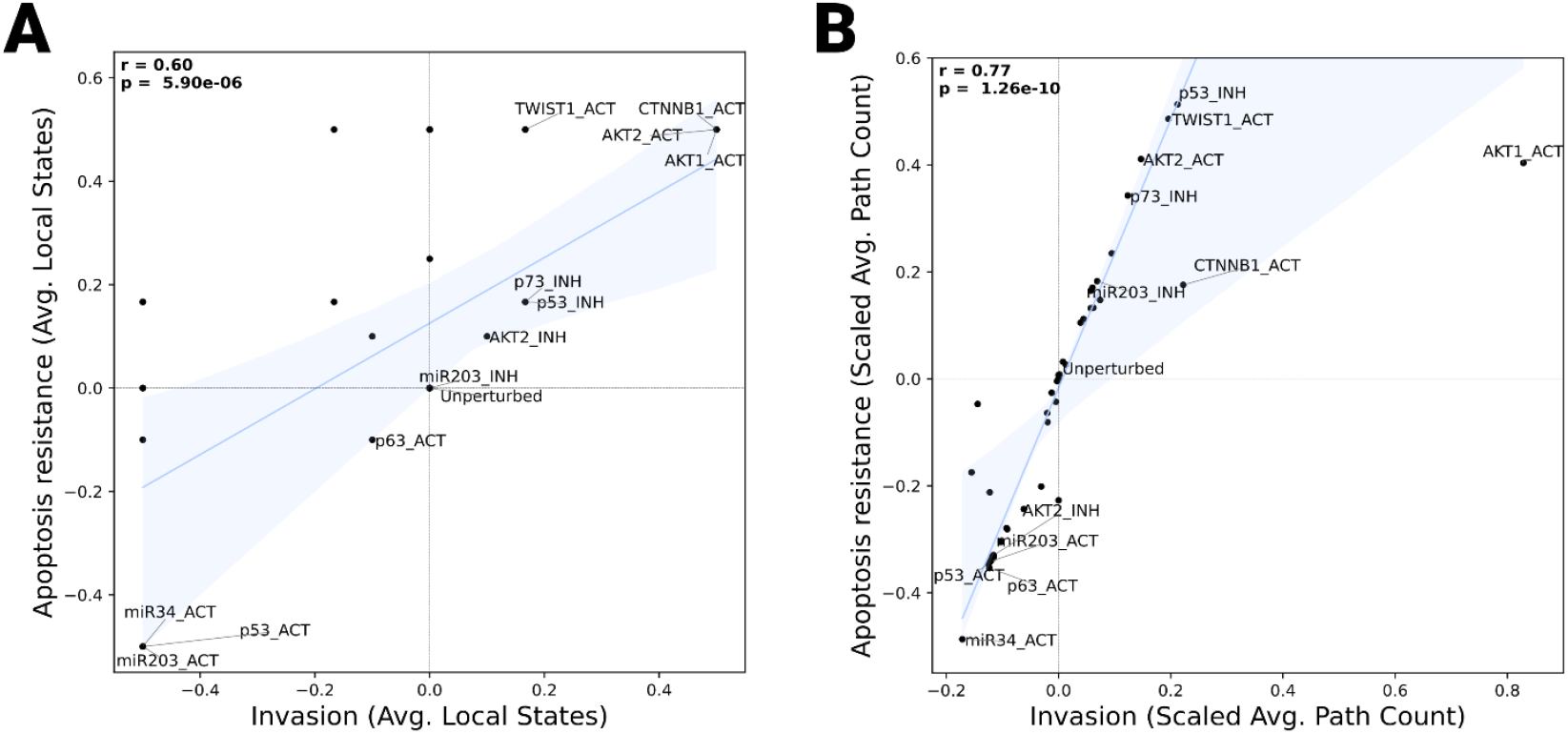
**A**. A scatter plot depicting the impact of additive perturbations on the Boolean states of the *Invasion* (X-axis) and Apoptosis (Y-axis) phenotype nodes. **B**. A scatter plot depicting the impact of additive perturbations on the path counts triggering the activation or inhibition of the *Invasion* (X-axis) and *Apoptosis* (Y-axis) phenotype nodes. The points represent *Invasion* and *Apoptosis* scores for each perturbation (n = 48) and the unperturbed setup, which are calculated as the average of *Invasion* and 1-*Apoptosis* local states (A) and path counts (B), min-max scaled (with reference at the unperturbed set up). Pearson correlation coefficient (r) and associated p-values are annotated in the upper-left corner of each plot. The regression line of best fit and estimate of 95% confidence interval are displayed in blue.

## Discussion

Boolean models, due to their simplicity and ability to represent cell signaling events, represent a powerful resource for studying the complex molecular dynamics of biological processes. However, their discrete nature limits their capacity to capture continuous aspects of cellular signaling, such as signal strength or the degree of activation or repression of biological entities (4). While some existing tools provide means to explore signal transduction, the computational demands of path exploration often force users to set limitations, such as maximum path length or computation time, to make calculations feasible (12, 19). As a result, a consistent workflow for assessing signal transduction strength in Boolean models is yet to be elucidated.

To fill this gap, we introduced BooLEVARD, an efficient tool for computing the number of paths leading either to node activation or repression. BooLEVARD allows the redefinition of node Boolean states to represent the set of non-redundant paths directly influencing the observed Boolean outcome. Additionally, BooLEVARD supports the introduction of node perturbations within a framework compatible with path calculations. To ensure interoperability with other tools, BooLEVARD reads Boolean models in the standard BoolNet format (21), and allows for the models to be exported back to the same format. Additionally, since the path collection computed by BooLEVARD is an intrinsic feature of Boolean models, the tool can be applied not only to analyze models representing biological processes but also to study signal strength in any Boolean model.

The usage of BooLEVARD in the metastatic model for the case study highlighted the importance and added value of focusing on signaling strength to better understand cell-fate decisions. Furthermore, the results were more precise and better aligned with biological findings, in comparison to SS analysis, as supported by the existing literature. The semi-continuous nature of BooLEVARD’s output enabled a more accurate identification of the most impactful perturbations, providing insights into which possible drug target nodes are likely to be more effective and which node mutations may be more strongly associated with metastatic risk, thus complementing current approaches toward the use of computational models in personalized medicine efforts (35).

### Availability and Future Directions

BooLEVARD is publicly available as a PyPi package at https://github.com/farinasm/boolevard, with user documentation detailing installation procedures and a step-by-step usage guide. The package documentation, including the API reference, was generated using Sphinx (36) to ensure clarity and ease of navigation, available at https://farinasm.github.io/boolevard/. To facilitate user onboarding, we provide Jupyter Notebooks alongside a toy example model, allowing users to explore BooLEVARD’s functionalities through a practical example. The source code used to generate the figures and supplementary figures presented in this paper is available at https://github.com/farinasm/boolevard/tree/main/paper.

Looking ahead, future versions of BooLEVARD will introduce features for identifying and incorporating positive feedback loops into the path count, further refining the tool’s ability to analyze signal transduction networks. Additionally, since simulations using Boolean models with larger networks (exceeding 100 nodes) or those with numerous stable states can be computationally demanding, upcoming updates will focus on optimizing performance to enhance scalability and efficiency.

## Author contributions

JZ, ET, and MF: Contextualization; MF: Design and Validation; ÅF and KL: Supervision; TA, ÅF, and KL: Funding Acquisition; all co-authors: Writing, review, and editing.

## Acknowledgements

We would like to thank the financial support from the Swedish Cancer Society (21 1888 Pj), and the Norwegian Cancer Society (216113) to KL, as well as the Novo Nordisk Foundation (NNF21OC0070381) to KL and TA. TA was supported by the Norwegian Cancer Society (grants 216104 and 273810), South-Eastern Norway Regional Health Authority (grants 2020026 and 2023105), and the Radium Hospital Foundation. ET and ÅF were supported by The Research Council of Norway (RCN) [grant number 310160], and under the framework of the European Research Area (ERA) PerMed program [grant number 329059]. Our gratitude also goes to Antonio Cabezas for his support with the code, and Professor Loïc Pauleve for his assistance with the conceptualization during the early stages of this work, Celia Montaña for the logo design, and Professor Martin Kuiper for his valuable insights and support.

